# A 3D-printed flow-cell for on-grid purification of electron microscopy samples directly from lysate

**DOI:** 10.1101/2023.03.17.533159

**Authors:** Kailash Ramlaul, Ziyi Feng, Caoimhe Canavan, Martín Natàlia de Garrido, David Carreño, Michael Crone, Kirsten E. Jensen, Bing Li, Harry Barnet, David T. Riglar, Paul S. Freemont, David Miller, Christopher H. S. Aylett

## Abstract

While recent advances in cryo-EM, coupled with single particle analysis, have the potential to allow structure determination in a near-native state from vanishingly few individual particles, this vision has yet to be realised in practise. Requirements for particle numbers that currently far exceed the theoretical lower limits, challenges with the practicalities of achieving high concentrations for difficult-to-produce samples, and inadequate sample-dependent imaging conditions, all result in significant bottlenecks preventing routine structure determination using cryo-EM. Therefore, considerable efforts are being made to circumvent these bottlenecks by developing affinity purification of samples on-grid; at once obviating the need to produce large amounts of protein, as well as more directly controlling the variable, and sample-dependent, process of grid preparation.

In this proof-of-concept study, we demonstrate a further practical step towards this paradigm, developing a 3D-printable flow-cell device to allow on-grid affinity purification from raw inputs such as whole cell lysates, using graphene oxide-based affinity grids. Our flow-cell device can be interfaced directly with routinely-used laboratory equipment such as liquid chromatographs, or peristaltic pumps, fitted with standard chromatographic (1/16”) connectors, and can be used to allow binding of samples to affinity grids in a controlled environment prior to the extensive washing required to remove impurities. Furthermore, by designing a device which can be 3D printed and coupled to routinely used laboratory equipment, we hope to increase the accessibility of the techniques presented herein to researchers working towards single-particle macromolecular structures.

## Introduction

Due to rapid and consistent advances in microscope hardware and image processing software, cryo-EM single particle analysis is able to determine macromolecular structures to well below 4 Å, in theory requiring only a tiny quantity of sample (as few as 10 000 particles) for a structure (Henderson, 1995). In practice, the sample and grid preparation stages of the workflow represent the major remaining bottlenecks to date (Elmlund & Elmlund, 2015; Thompson *et al*, 2016; Arnold *et al*, 2018). Typically, recombinantly produced proteins and complexes are purified to homogeneity and concentrated to between 0.1–0.5 mg/mL. This is a dramatically lower requirement than that for X-ray crystallography, the structure determination method which holds the current record for number of structures as well as highest attained resolution. Yet, due to the methods by which grid preparation is routinely performed, more than 99% of the sample solution applied is lost during the blotting process required to yield a thin film during grid preparation and vitrification (Kemmerling *et al*, 2012; Arnold *et al*, 2018).

A significant problem, which is exacerbated by this huge loss of sample during grid preparation, is that samples are often precious, by virtue of being extremely difficult to make, requiring large quantities of expensive tissue culture reagents. This is especially true for the production of large, often mammalian, macromolecular complexes for which cryo-EM is the most practical method that can be used to perform structural studies, and for which expensive mammalian tissue culture systems are the only suitable path to recombinant expression. Throughout the typical purification approach, with multiple rounds of chromatography aiming to retain sample and discard impurities, the sample is slowly obtained in a purer form. For example, during immobilised metal affinity chromatography, polyhistidine-tagged sample proteins are bound by the affinity matrix and buffers containing low levels of imidazole are used to wash away nonspecifically-bound contamination. However, the overall process is cumbersome: it can take days to complete, the sample yield steadily decreases with each chromatographic step, and the purified samples (especially macromolecular complexes) are often not stable for long periods (Waugh, 2005; Pina *et al*, 2014). Thus, an innovative solution to extracting as much as possible from raw inputs without the unnecessary wastage of current workflows could be to perform single-step purification steps on TEM grids themselves, using an affinity tag system.

It is already common practice for electron microscopers to add additional support films to TEM grids, as this helps achieve a crucial condition for imaging: that samples must be vitrified within ice thin enough to ensure high contrast in micrographs, whilst retaining the structural integrity of the sample itself sufficiently to reflect the biologically active state (Glaeser, 2018). Thin vitreous ice increases the exposure of particles to the air-water interface (Noble *et al*, 2018). At this interface, biomolecules are subjected to harsh conditions which often lead to denaturation and preferential orientation (Han *et al*, 2017; Glaeser, 2018; D’Imprima *et al*, 2019). The most commonly used support films are amorphous carbon, graphene oxide and graphene; the latter two being preferable due to their electron transparency relative to amorphous carbon, and a suite of methods are available which allow their reproducible addition to TEM grids in EM workflows (e.g. de Martín Garrido *et al*, 2021). A key advantage to the use of an additional support film is that sample particles immobilised in such a way can be protected from the air-water interface, and its associated refractory effects, by virtue of being held at the opposite face of the liquid film (D’Imprima *et al*, 2019). Indeed, it has been found that support films are protective for biomolecules and can also increase their density and retention on grids, aiding downstream data collection and image analysis (Pantelic *et al*, 2010; Russo & Passmore, 2014; Glaeser, 2018). Thus, support films are uniquely positioned to synergistically tackle both the sample-and grid-preparatory stages of cryo-EM workflows: by performing affinity capture on the grid surface, it should be possible to reduce both causes of sample loss whilst maintaining sample integrity for imaging.

There is a large body of work showing that on-grid affinity capture methods are practicable. A seminal proof-of-concept study in the early 2000s showed that 2D streptavidin crystals deposited onto a TEM grid (as first produced by Darst *et al*, 1991) could be used to immobilise biotinylated DNA substrates and a RNAPI-oligonucleotide complex to the surface of a holey foil grid (Crucifix *et al*, 2004). This method was then applied to tethering of membrane protein-containing liposomes, functionalised with biotinylated lipid headgroups, to 2D streptavidin crystals grown directly on a TEM grid (Wang *et al*, 2008). These authors showed that only a small amount of proteoliposome is required due to enrichment on the grid surface through affinity interaction. Another group reported direct binding of chemically biotinylated proteins to 2D streptavidin crystals using input protein concentrations as low as 10 µg/mL (Han *et al*, 2012), and optimisation of this 2D crystal-based affinity grid culminated in the structure of the *E. coli* ribosome to 3.9 Å resolution, after subtraction of the streptavidin signal (Han *et al*, 2016). Additionally, use of streptavidin-based affinity grids reduced preferential orientation of the yeast RNAP elongation complex (Lahiri *et al*, 2019). It has also been demonstrated, by another group, that chemical functionalisation of amorphous carbon and derivatisation with Ni-NTA and Protein G provides another variety of affinity support, capturing His-tagged valosin-containing protein from samples at concentrations as low as 0.1 nM (Llaguno *et al*, 2014). Amorphous carbon, however, contributes significantly to background scattering, and can impede higher resolution reconstruction.

The principle of on-grid affinity purification, namely the use of a derivatised grid surface to specifically enrich a tagged sample on its surface from a mixture of sample components, was first formalised by Kelly and colleagues, with what they termed ‘monolayer purification’ (Kelly *et al*, 2008b). These authors used lipid monolayers functionalised with Ni-NTA to specifically bind His-tagged proteins, which they showed could be enriched from as little as 0.4 µg/mL in cell extracts. Their method involved the generation of a functionalised lipid monolayer on top of a droplet of cell extract containing, or recombinantly expressed and purified, His-tagged protein sample in a Teflon block. After sufficient time had passed to allow for affinity capture of the sample, a TEM grid was placed onto the monolayer to transfer the sample to the grid (Kelly *et al*, 2008b). The principle of this method was then expanded to make the grids more robust (Kelly *et al*, 2008a), as well as to modify functionalised lipid monolayers with antibodies to capture more specific, non-tagged targets from cell extracts (Kelly *et al*, 2010) or to remove the need for monolayers entirely (Yu *et al*, 2014). On the basis of their previous work, Yu and colleagues determined the first high-resolution structure of the Tulane virus to 2.6 Å resolution, fabricating anti-Tulane virus IgG-coated TEM grids and capturing virus particles directly from cell extracts (Yu *et al*, 2016b, 2016a). Whilst the resolution of structures determined in these studies steadily increased, lipid monolayer-and amorphous carbon-based affinity grids still suffer strongly from affinity matrix instability, significant background scattering and additionally the need to optimise each sample empirically.

As mentioned above, graphene oxide and graphene are touted to be the best sample support films for cryo-EM work, given their electron transparency, thickness, and stability to thermal motion in comparison to amorphous carbon (Pantelic *et al*, 2011). In particular, graphene oxide (GrOx) has been the subject of several affinity grid studies, given its relatively diverse range of reactive functional groups compared to graphene. Benjamin and colleagues functionalised GrOx flakes with Ni-NTA to capture His-tagged proteins, both pre-purified and from cell lysates, and showed that their grids are mechanically superior to lipid monolayer-based affinity grids (Benjamin *et al*, 2016). The process of derivatisation, however, is intensively laborious and requires specialist equipment which is likely not to be readily available to many structural biology research groups. A significant improvement to the modification of graphene oxide was made by Wang and colleagues, who functionalised their graphene oxide-supported TEM grids using Cu-free click chemistry (Wang *et al*, 2020). These authors derivatised their grids using either SpyCatcher or its cognate binding partner SpyTag, a split-domain tag-partner pair capable of forming a spontaneous isopeptide bond between two key residues, capable of rendering tag binding irreversible (Zakeri *et al*, 2012). This method required two, at minimum, heterobifunctional crosslinking reagents containing large polyethyleneglycol (PEG) linkers, proving far less laborious and costly. The authors spiked rabbit reticulocyte lysate with a known amount of SpyTagged apoferritin and demonstrated successful affinity capture and removal of nonspecific background; however, the affinity purification of a heterologously expressed protein direct from native sources remained to be shown.

In this study, we have demonstrated and make widely available a simple-to-use and readily practical flow-cell device for on-grid affinity purification of EM samples. Our method allows on-grid affinity purification through the extensive application of inputs such as whole cell lysate containing the particle of interest at an arbitrary concentration, and enables equally extensive washing of the resulting grids, which is necessary to remove impurities, taking advantage of the spontaneous covalent bond formation between SpyCatcher003, which we used to functionalise the surface of graphene oxide-based TEM grids, and its cognate partner SpyTag003. To the best of our knowledge, this is the first proof-of-concept of a flow-cell capable of such extensive application and washing. Our solutions are designed to be as simple and as readily adopted as possible; our flow-cell can be 3D-printed either in-house with relatively inexpensive materials and equipment, or ordered from commercial providers, and uses standard attachments that can be readily coupled to widely available liquid chromatography equipment but can be run with as little as only a peristaltic pump if required. Our flow-cell design and setup allows the generation of reliable, consistently populated, readily washed, graphene oxide-based affinity grids suitable for TEM studies.

## Results

### Simplification of the production and storage of graphene oxide affinity grids using Cu-free click chemistry by optimisation of the method of Wang and colleagues

To prepare graphene oxide-based affinity grids, we simplified and improved upon the protocol of Wang and colleagues, which utilised SpyCatcher derivatisation of synthesised GrOx sheets by Cu-free click chemistry (Wang *et al*, 2020). Instead of synthesising graphene oxide, we instead sought to improve the accessibility of this protocol by using commercially available graphene oxide flakes, which can be purchased as dispersions in water and are sold by major biochemical reagent vendors (such as Merck) (Fig. 1a). We applied GrOx support films to TEM grids using previously described methods (Cheung *et al*, 2018; de Martín Garrido *et al*, 2021b), and showed that, although a low level of GrOx fragmentation and agglomeration was observed when comparing grids before (Fig. 1b, left) and after (Fig. 1c, left) washing, overall the GrOx support films withstood the functionalisation steps and rigorous washing procedures and a large proportion of holes per grid square remained suitable for imaging (Fig. 1c, right).

**Figure 1:**
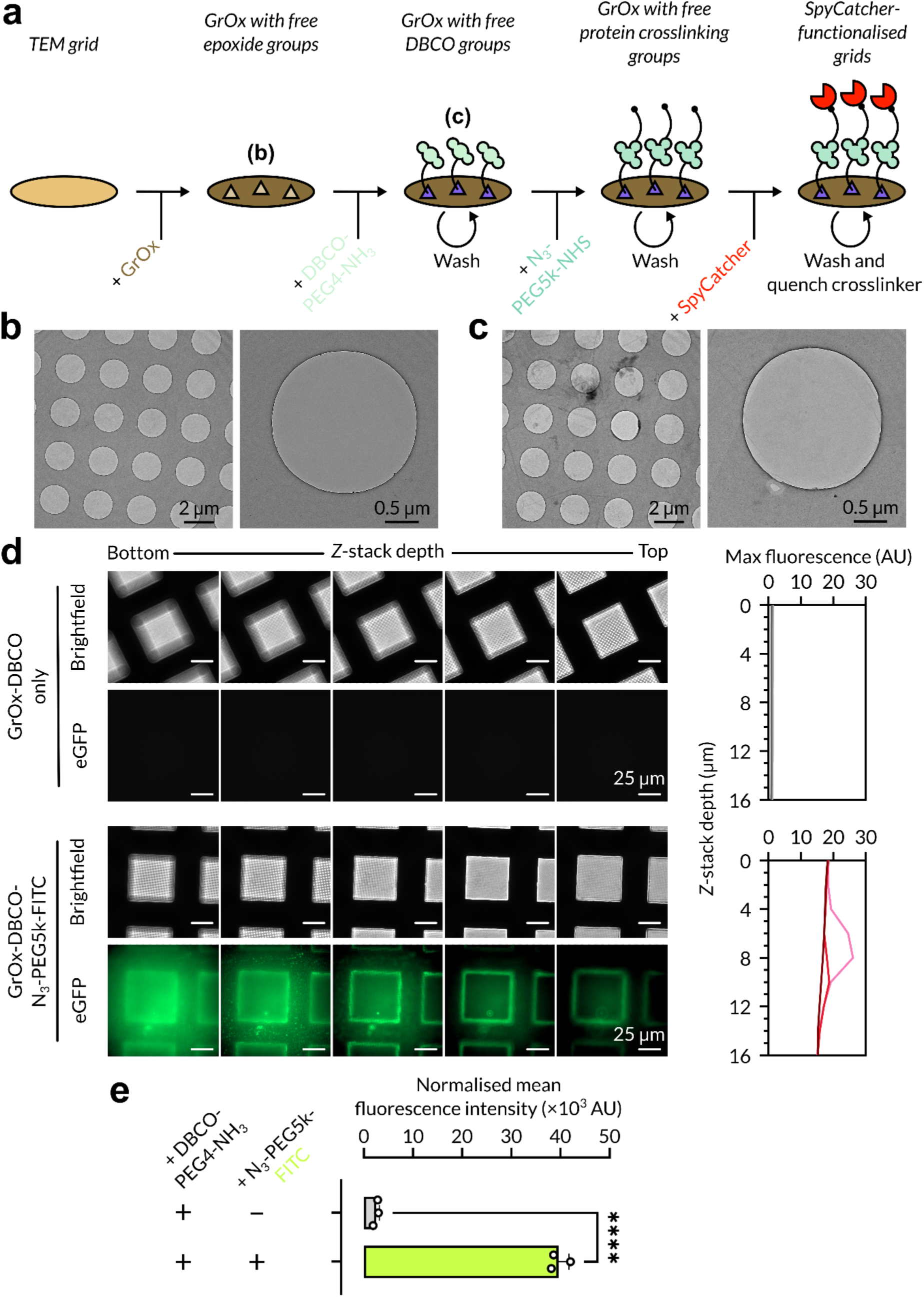
Preparation and functionalisation of graphene oxide-based affinity grids. **a.** Schematic overview of graphene oxide functionalisation and derivatisation based on established protocols (Wang *et al*, 2020). **b.** Grid square and representative hole image with visible GrOx coverage after GrOx support film addition. **c.** Grid square and representative hole image with visible GrOx coverage after complete functionalisation of grids as in a, showing increased GrOx agglomeration after washing process, but retention of pristine areas for imaging. **d.** *Z*-stacks with brightfield and eGFP illumination of DBCO-only grid square view (top) and DBCO-azide-FITC-functionalised grid square view (bottom). Normalised maximum fluorescence values plotted by *Z*-stack depth is shown next to each condition. **e.** Normalised mean fluorescence intensity comparison between GrOx-DBCO-only grid and DBCO-azide-FITC-functionalised grid. Error bars represent standard error of the mean (*n* = 3). \*\*\*\**P* < 0.0001.

We then proceeded to derivatise our GrOx affinity grids using Cu-free click chemistry between DBCO-PEG4-amine and azide-PEG5k-crosslinker compounds (Wang *et al*, 2020). We first validated the functionalisation of commercial GrOx flakes with DBCO-PEG4-amine using the fluorescent molecule azide-PEG5k-fluorescein isothiocyanate (FITC). We used fluorescence microscopy to compare the affinity grids by collecting Z-stacks vertically at the grid square level (see Materials & Methods). Whereas we observed very low signal in the DBCO-only grid, which likely represents autofluorescence of the GrOx flakes as compared to GrOx-coated grids without functionalisation, the azide-PEG5k-FITC-derivatised grid showed significant fluorescent signal which was distributed across the illuminable area of the grid square (Fig. 1d). Mean fluorescence intensity (MFI) normalised to grid square area showed a 15-fold increase in fluorescence intensity of the FITC-derivatised grid in comparison to the DBCO-only control (Fig. 1e). Additionally, the Z-stack also allowed identification of the approximate vertical displacement of the maximum fluorescent output, which would likely correspond to the grid surface; utilising the fluorescent maxima would also serve to control for the gradient of normalised MFI values caused by GrOx autofluorescence. As expected, azide-PEG5k-FITC-derivatised grids showed fluorescence maxima at distinct depths through the z-stack, whereas the DBCO-only grids did not produce any comparable signal nor significant maxima (Fig. 1d, plots). Taken together, this data indicated that commercially available GrOx flakes were suitable for affinity grid production in place of GrOx produced by Wang and colleagues’ modified Hummer’s synthesis (Wang *et al*, 2020).

We chose to functionalise our affinity grids with N-hydroxy succinimide (NHS) ester functional groups, which stably crosslinks to free amine groups e.g. lysine sidechains, in DMSO rather than water, to prevent passive hydrolysis and extend shelf life after production as DMSO is an aprotic solvent. We tested this crosslinking using purified mGreenLantern, a bright GFP variant (Campbell *et al*, 2020), which we incubated on affinity grids derivatised with azide-PEG5k-NHS ester. In the absence of mGreenLantern, we observed background fluorescence comparable to GrOx autofluorescence from previous experiments when NHS ester-affinity grids were subjected only to Tris buffer quenching after derivatisation (Fig. 2a). We did observe some nonspecific binding of mGreenLantern to GrOx-DBCO only grids, which manifested as bright punctae associated with high fluorescence maxima (Fig. 2b). However, as expected, when we added mGreenLantern to NHS ester-derivatised grids we observed fluorescence coverage across the complete area of each affinity grid (Fig. 2c), corresponding to a ∼7-fold greater specific fluorescence (Fig. 2c & d) demonstrating successful affinity binding. The observation of bright punctae in both the GrOx-DBCO only and NHS ester-derivatised grid conditions suggests that these features represent nonspecifically-interacting aggregates of mGreenLantern.

**Figure 2:**
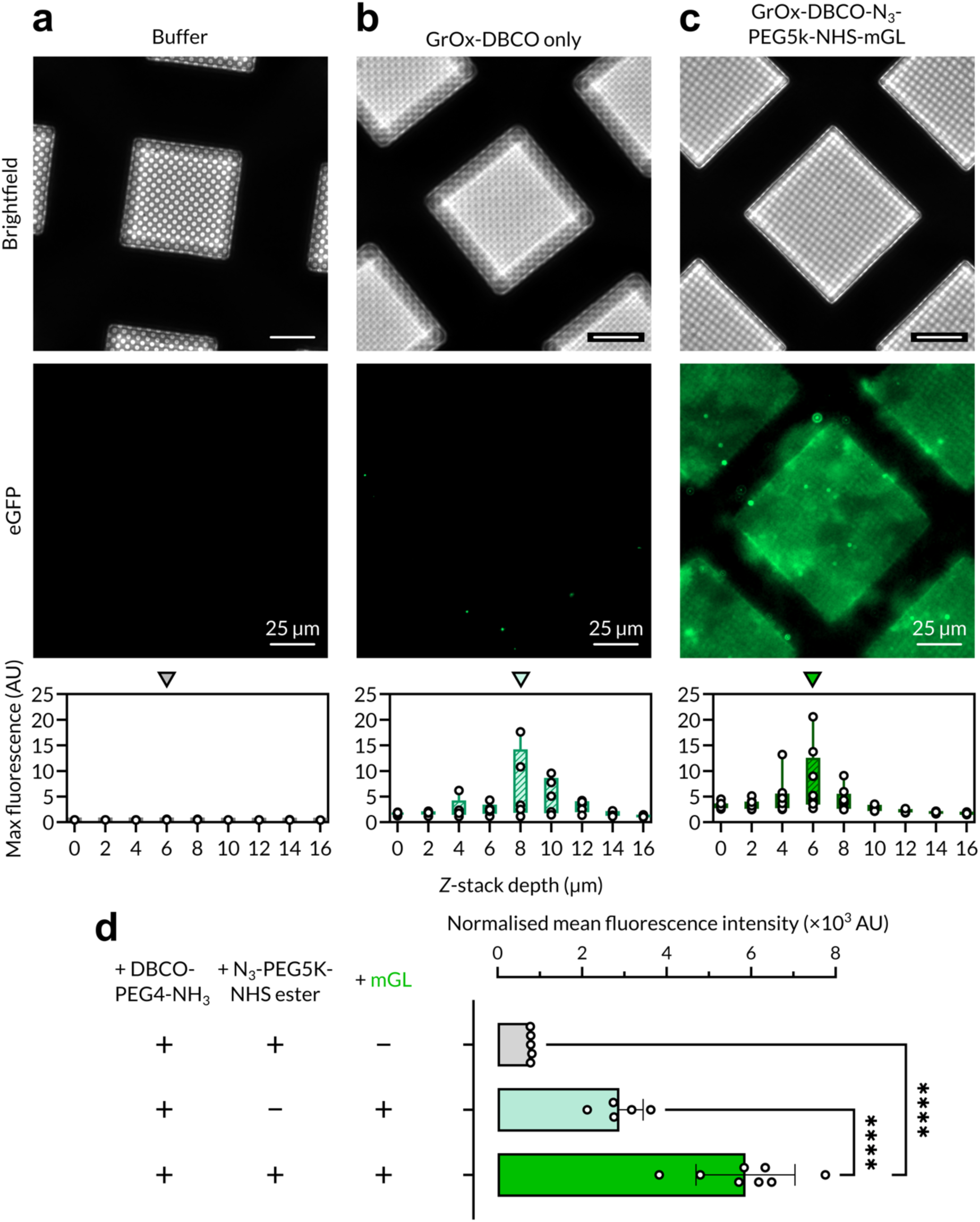
Validation of amine-NHS ester-based crosslinking for derivatisation. **a.** Brightfield (top), eGFP (bottom) channel images and normalised maximum fluorescence plots for DBCO-azide-NHS ester-functionalised affinity grids with buffer only **(a)**, DBCO-only grids with mGreenLantern **(b)** and DBCO-azide-NHS ester-functionalised affinity grids with mGreenLantern **(c)**. Arrows above maximum fluorescence plots indicate *Z*-stack depth shown in images, corresponding to slices with maximum fluorescence intensity. **d.** Normalised mean fluorescence intensity comparison betweenDBCO-azide-NHS ester-functionalised affinity grids with buffer only, DBCO-only grids with mGreenLantern and DBCO-azide-NHS ester-functionalised affinity grids with mGreenLantern (mGL). Error bars represent standard error of the mean. \*\*\*\**P* < 0.0001.

To expedite the process of fabricating affinity grids, and simultaneously reduce the complexity of the manual handling steps required for transferring grids between reagents, we designed a 96-well plate format tool (Fig. 3a). The plate conforms to standard 96-well spacing (8.99 mm separation) along the long axis, however the separation between wells was increased to 9.6 mm along the shorter axis to allow use of apparatus holding eight standard tweezers side-by-side simultaneously but retain compatibility with 96-well dispensing apparatus. The main advantage of using the tool was that by creating wells with slots situated within broader, tapered cavities, we could place grids vertically to remain accessible by tweezing, thus performing the incubation steps required in the 96-well plate rather than using individual Eppendorf tubes for each grid (Fig. 3b). We optimised this tool for fabrication by computer numerical control (CNC) milling to allow the use of polytetrafluoroethylene (PTFE) as a material for the plate, which is more compatible with organic solvents, and used it repeatedly to fabricate our affinity grids.

**Figure 3:**
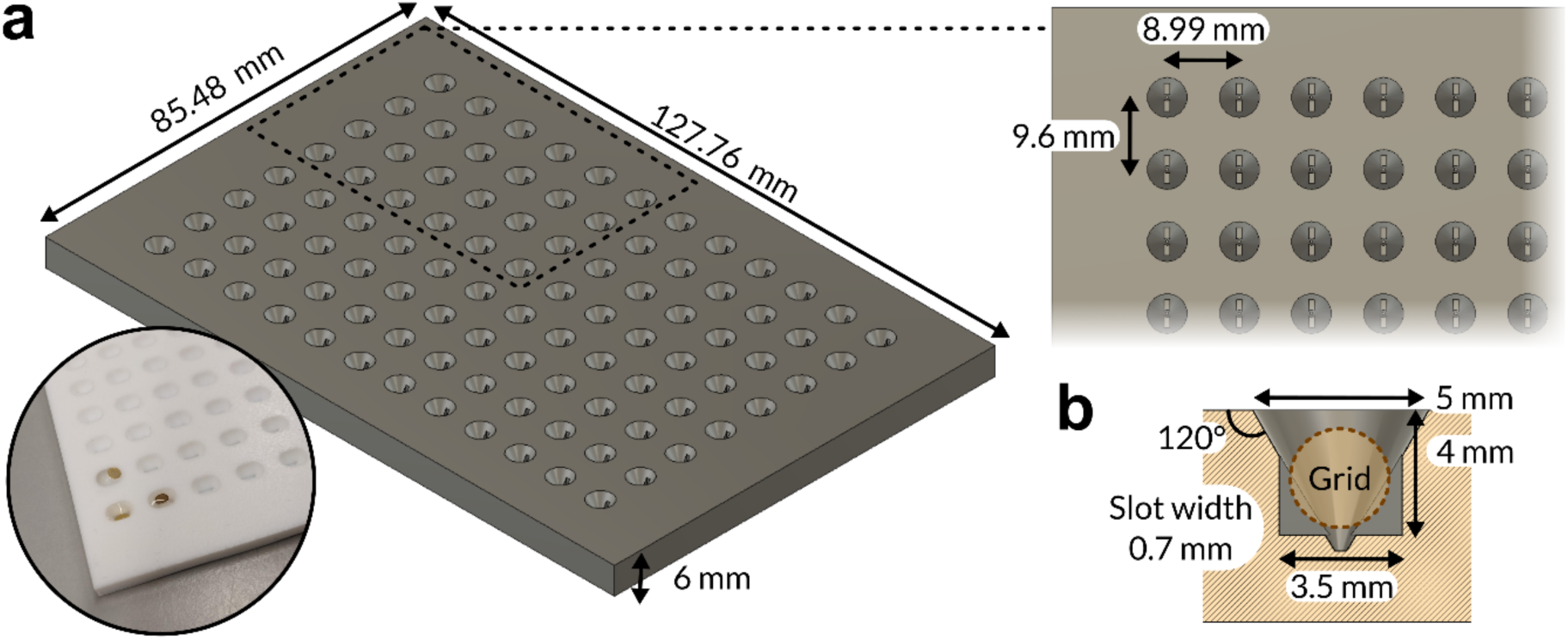
Design of a 96-well plate tool for efficient affinity grid derivitisation. **a.** Schematic of the 96-well affinity grid tool and top-down view of individual well. The inset image shows a picture taken of the plate tool with grids either submerged within or floating atop buffer placed in wells. **b.** Side-view section of individual well.

### A novel 3D-printable flow-cell enables application of tiny (100s of uL) to substantial (100s of mL) volumes over affinity grids and their subsequent recovery as electron microscopy samples

On-grid affinity purification of tagged samples from heterogeneous mixtures (e.g. lysate) is already practicable if the tagged sample is present in high concentrations and the present contaminants do not strongly non-specifically adhere to the surface of the grid. Typically, however, these requirements are not met, therefore both extensive application of sample to the grid sample to bind sample, and rigorous washing of the surface to remove bound contaminants, are needed. To achieve this, we designed a new kind of sample support block (de Martín Garrido *et al*, 2021a) (Fig. 4a), which is 3D-printable, i.e. using stereolithography printers, so as to make this method more accessible to labs which may not possess more complicated and bespoke microfluidic setups as purchasing 3D printed parts is vastly cheaper and easier.

**Figure 4:**
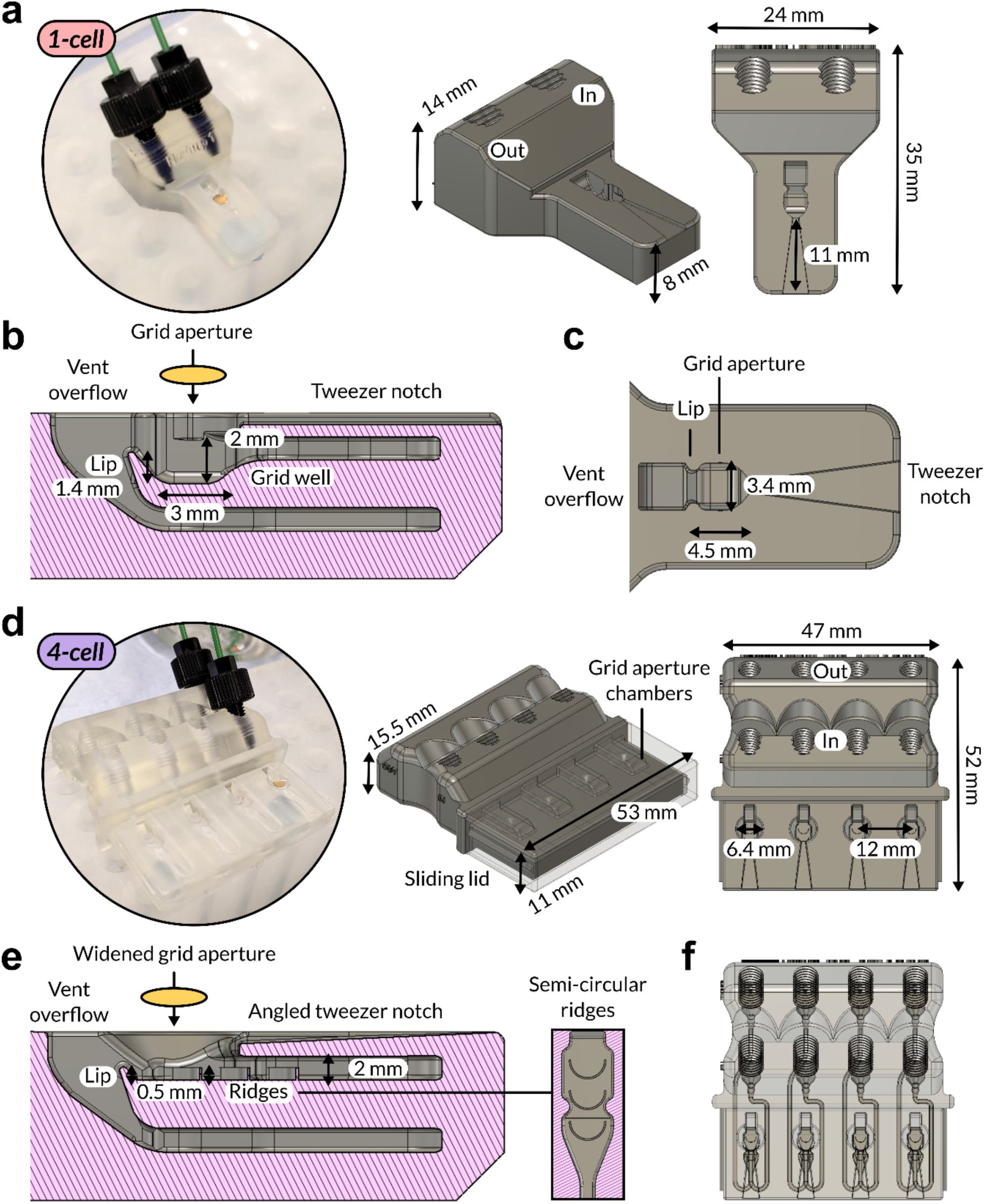
Design of 3D-printable flow-cell devices for on-grid affinity purification workflows. **a.** Picture and schematic of the flow (1)-cell device design from two different angles, showing key device dimensions. Inlet and outlet ports are labelled. The inset image shows a picture taken of the 1-cell during operation. **b.** Side-view section of 1-cell flowpath through grid aperture. A TEM grid is indicated by the orange oval. **c.** TEM grid aperture design, showing aperture where grids are placed using the tweezer notch, and lip and vent overflow where buffer pumped into the flow-cell flows into and can then be pumped out of the device. **d.** Picture and schematic of the flow (4)-cell device design from two different angles, showing key device dimensions. A sliding lid designed to cover the grid apertures is shown as semi-transparent. The inset image shows a picture taken of the 4-cell during operation. **e.** Side-view section of 4-cell flowpath through grid aperture. A TEM grid is indicated by the orange oval. Key differences to the 1-cell design include a wider grid aperture, angled tweezer notch, a flat inlet channel and the addition of semi-circular ridges along flowpath, also shown top-down as inset, to induce turbulent flow under the grid surface. **f.** Semi-transparent render of 4-cell, showing independent inlet and outlet flowpaths.

The flow-cell (‘1-cell’) features inlet and outlet screw ports to allow attachment to laboratory equipment such as peristaltic pumps and ÄKTA purification systems which utilise standardised (e.g. 1/16”) union fittings and tubing diameters (Fig. 4a). The inlet and outlet ports channels run through a flow cell which consists of vertically displaced flowpaths separated by a lip over which buffer flows and is drawn away (Fig. 4b), with a grid aperture surrounded by baffles in the inlet flowpath, into which a TEM grid can be held with tweezers (Fig. 4c). The U-bend mechanism (Hume & Ives, 2009) inspired the design of the lip (Fig. 4b, lip), which allows the maintenance of essentially constant liquid levels while the block is in operation. To operate the flow-cell, we used a peristaltic pump configured with two tubing circuits; input, drawing buffer to the inlet screw port, and output, drawing buffer from the outlet screw port into a waste container. The flow-cell device worked optimally by adjusting the output circuit to run slightly faster than the input, avoiding buffer flow outwards through the grid aperture during operation.

We also designed a version of the flow-cell optimised further for cryo-EM studies, which could support the preparation of up to 4 affinity grids in parallel (Fig. 4d), useful for cryo-EM as in this case grids are typically prepared in batches of 4. This ‘4-cell’ version contained the basic design features of the 1-cell, with some modifications made after empirical optimisation. These key changes include: a wider grid aperture (Fig. 4d, 6.4 mm diameter aperture); an angled tweezer notch, to discourage capillary flow upwards along the tweezer groove; a flat inlet channel; and, the addition of semi-circular ridges along each inlet flowpath (Fig. 4e). Semi-circular ridges have been determined to act as efficient passive micromixers which encourage turbulent flow (Okuducu & Aral, 2019). We placed these ridge elements in the flow path entering and under the grid aperture to increase the efficiency of washing (Fig. 4e, inset). Liquid flow is considered to be highly laminar and variably viscous with low molecular diffusion constants — thus, when washing the grid surface, it is likely that only the uppermost ‘surface’ of the liquid contacts the grid. Computer simulations using semi-circular ridges suggest that the ridges cause helicoidal flow patterns (Okuducu & Aral, 2019), which — for laminar flow in a straight channel — would generate fluid flow across the grid perpendicular to the direction of mass flow, and thus improve washing efficiency. The ridges clearly disrupted laminar flow and increased the amount of turbulence when buffer flowed through the grid aperture section, observed using dye and buffer. Finally, we arranged 4 completely independent inlets and outlets side-by-side in our 4-cell device to overcome prior challenges in parallelising flow (Fig. 4f).

### Using our apparatus, MS2-SpyTag003 virions could be efficiently captured on affinity grids directly from cell lysate, and off-target contaminants could be effectively washed away

To test the utility of the affinity grid support block, we cloned the viral MS2 coat protein (CP) with the SpyTag003 (Keeble *et al*, 2019) sequence as a dimer (CP-SpyTag003-CP). This strategy was devised as MS2 virion production was optimised for another study conducted in our laboratory (de Martín Garrido *et al*, 2020) and could provide a useful test sample easily observable directly in EM micrographs, and trivially and accurately countable for the purposes of quantification and statistical validation, as opposed to indirectly as with fluorescence measurements. The construct design was inspired by a study investigating MS2 morphology, specifically their ‘SpyTag7’ construct (Biela *et al*, 2022). As a preliminary test, semi-purified MS2 virions with the CP-SpyTag003-CP (MS2-SpyTag003) were crosslinked *in vitro* to purified SpyCatcher003, using the shift of the CP band to higher molecular weight after SDS-PAGE as evidence of crosslinking (Fig. 5a). We indeed detected a steady increase in two higher-MW bands. Both of these gel bands were confirmed by mass spectrometry (MS) to contain MS2 CP, thus suggesting that MS2-SpyTag003 was a suitable sample to test affinity capture.

**Figure 5:**
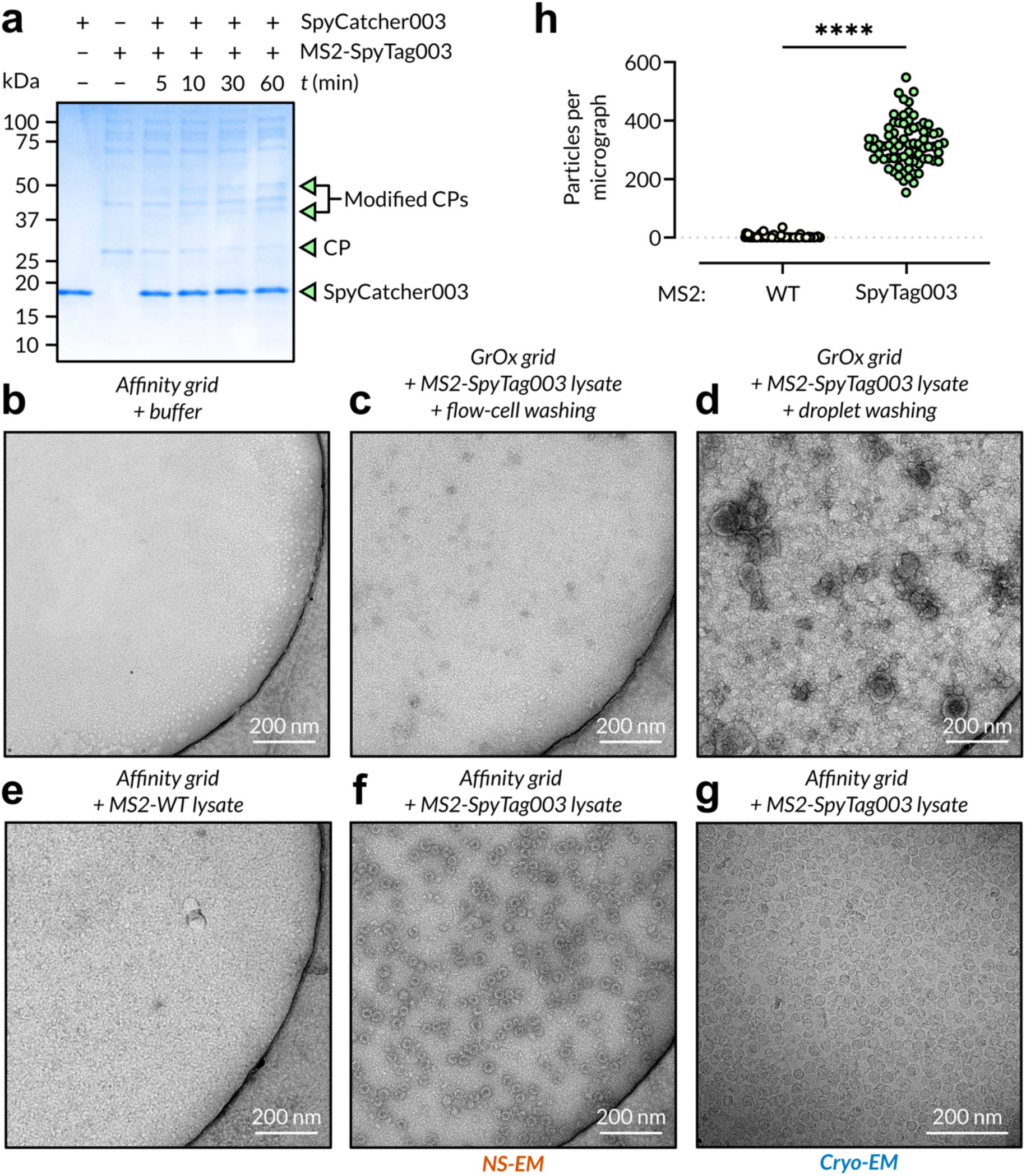
Use of our flow-cell devices enabled direct capture of SpyTag-MS2 virions from lysate and extensive washing resulted in the retention of only insignificant levels of contamination. **a.** *In vitro* crosslinking assay between semi-purified MS2-SpyTag003 and purified SpyCatcher003. Characteristic bands for SpyCatcher003 and MS2 coat protein (CP) are labelled. Modified CPs were confirmed by MS. **b.** Negatively-stained micrograph of SpyCatcher003-functionalised NHS ester-affinity grid washed with buffer using the flow-cell device. **c.** Negatively-stained micrograph of SpyCatcher003-functionalised NHS ester-affinity grid loaded with MS2-SpyTag003 lysate and subsequently washed with buffer using the flow-cell device. **d.** Negatively-stained micrograph of SpyCatcher003-functionalised NHS ester-affinity grid loaded with MS2-SpyTag003 lysate and subsequently washed using sequential buffer droplets. **e.** Negatively-stained micrograph of SpyCatcher003-functionalised NHS ester-affinity grid loaded with MS2-WT lysate and subsequently washed with buffer using the flow-cell device. f. Negatively-stained micrograph of SpyCatcher003-functionalised NHS ester-affinity grid loaded with MS2-SpyTag003 lysate and subsequently washed with buffer using the flow-cell device. **g.** Cryo-electron micrograph of affinity grid loaded with MS2-SpyTag003 lysate as in **f. h.** Quantification of MS2 particles per NS-EM micrograph from triplicate affinity grids loaded with either MS2-WT or MS2-SpyTag003, showing a statistically significant difference as compared by unpaired *t*-test (*n* = 75). \*\*\*\**P* < 0.0001.

We then sought to validate our affinity grid and purification system using this sample. We functionalised NHS ester affinity grids with SpyCatcher003 and tested several conditions using negative-staining EM (NS-EM). First, we imaged a functionalised grid which had been washed with flow-cell washing buffer to observe the expected background; although some negative-staining artefacts were present around the edges of the holes in these grids, there was no detectable sample (Fig. 5b). We then tested the capacity of the flow-cell for washing MS2-SpyTag003 lysate from the surface of regular, non-functionalised GrOx grids. These grids were extremely similar to the buffer-only grids, showing a very slight increase in contamination (Fig. 5c), suggesting that the flow-cell was necessary to achieve wash-out of contamination but sufficient to do so. Conversely, GrOx-only grids incubated with lysate and washed using up to 5 sequential 30 µL buffer droplets stained very poorly due to extremely high levels of contamination (Fig. 5d). We also wanted to determine whether affinity grids could be similarly treated to wash and remove nonspecific samples and contamination; using MS2-WT lysate, we found again a low-level increase in background contamination compared to the buffer-only grids (Fig. 5e). Lastly, we tested for specific affinity capture of MS2-SpyTag003 by functionalised affinity grids, and observed a high concentration of well-distributed particles matching known MS2 morphology using both NS-EM (Fig. 5f) and cryo-EM (Fig. 5g). We compared specific affinity capture of MS2-SpyTag003 versus MS2-WT by collecting NS-EM datasets of each across 3 independently prepared grids. After particle selection, using the well-known morphology of MS2, we observed significant capture of MS2-SpyTag003 versus MS2-WT, indicating that affinity purification was highly specific (Fig. 5h).

## Discussion

To successfully retrieve samples of interest, such as protein complexes, from a heterogeneous input such as cell lysate, it is necessary to ensure efficient capture of proteins of interest at the grid surface in a controlled environment, whilst simultaneously supporting extensive washing of TEM grids thereafter. To do so, we required a robust affinity grid with a passivated surface and a device which could physically house our grids. This device would provide a system by which to flow sample, containing tagged macromolecular complexes of interest, over the functionalised surface of the grid, without incurring sample binding to the other side, which would cause greater background scattering. Additionally, given the requirement to flow sample over the affinity grid, it would then also be possible to extensively wash the grid surface to remove impurities and contamination, such as is performed during standard chromatographic protein purification workflows. We built upon prior work in our lab (de Martín Garrido *et al*, 2021a) and designed our flow-cell to be 3D-printable using SLA printers, so as to make this method more accessible to labs which may not possess more complicated and bespoke microfluidic setups, as purchasing 3D printed parts is vastly cheaper and easier.

We adapted the protocol of Wang and colleagues who used Cu-free click chemistry to functionalise the surface of synthesised GrOx with reactive sulfhydryl groups (Wang *et al*, 2020). We sought to validate this method using commercially available GrOx flakes, which would increase the accessibility of the on-grid affinity purification approach, finding that GrOx flakes tolerated numerous washing steps and functionalisation proceeded efficiently. DBCO-PEG4-amine and azide-PEG5k-crosslinker reagents were prepared in DMSO, as it is an aprotic solvent which protected the reactive groups from passively degrading. We were able to use derivatised grids for functionalisation after months if stored in DMSO at room temperature. Finally, we chose to use NHS ester-based crosslinking chemistry, as it would also be straightforward to directly crosslink purified protein samples to NHS ester-derivatised grids. This could be a worthwhile grid preparation strategy for complexes which have established purification strategies but are very low-yield. Extensive application of such dilute samples could be facilitated through the use of one of our flow cells to ensure a sufficiency of the target complex coming into contact with the derivatised surface.

Our 1-cell design was conceived to provide a proof-of-principle test of the on-grid affinity purification workflow, validated by NS-EM using SpyTagged MS2 virions. MS2-SpyTag003 represented an ideal sample due to its size and distinct morphology in electron micrographs, making it a useful target for quantification. Our initial testing confirmed that the anticipated extensive washing was required to remove impurities after the application of cell-lysate to grids, droplet washing proving insufficient to provide a clean field of view for imaging (Figure 4). Moving to affinity capture, we found that we could achieve optimal coverage by incubating affinity grids on SpyCatcher003 solution for 30 mins; for samples of widely different sizes, however, coverage will need to be optimised on a sample-by-sample basis. It was clear from our MS2-SpyTag003 experiments, however, that the on-grid affinity purification approach using our novel flow-cell device should significantly decrease the amount of input sample required. For example, we can postulate that under optimal circumstances for a 3 mm TEM grid whose ideal surface coverage would be 20% for single particle analysis, and using MS2-SpyTag003 virus with a 30 nm particle diameter and approximate titre of ∼10^9^ particles/mL, we would require only 2.1 mL of MS2-SpyTag003 lysate to obtain satisfactory surface coverage. In practise we found that we were able to perform on-grid affinity purification experiments using as little as 125 µL of neat MS2 lysate per affinity grid while achieving an observed coverage of ∼8.6% on average based upon mean MS2-SpyTag003 count (Fig. 5f, g). This reduction in input requirements to several millilitres or fewer of neat lysate is an improvement on the order of hundreds of fold over the general expression volume requirements for chromatographic protein purifications (Pina *et al*, 2014). We envisage that the on-grid affinity purification method with the flow-cell device presented herein can begin to address the sample-and grid-preparation stages of cryo-EM workflows, in particular for difficult-to-produce, low-yield macromolecular samples.

## Materials and Methods

### Recombinant protein cloning, expression & purification

SpyCatcher003 (Keeble *et al*, 2019) and mGreenLantern (Campbell *et al*, 2020) were ordered as gBlock fragments (IDT) codon-optimised for *E. coli* expression. SpyCatcher003 was cloned into the expression vector pET9a with a TEV cleavage site followed by a His6-tag directly in-frame at the C-terminus (pET9a-SpyCatcher003-TEV-His6). mGreenLantern was cloned into the pET9a vector with a (GGS)9 linker immediately between the mGreenLantern ORF and TEV cleavage site (pET9a-mGreenLantern-(GGS)9-TEV-His6). Plasmids prepared of each construct were validated by sequencing (Eurofins Genomics).

For both SpyCatcher003 and mGreenLantern, expression and purification was performed identically. Plasmids were transformed into *E. coli* strain BL21(DE3) for recombinant protein expression. 1 L LB supplemented with 50 µg/mL kanamycin was inoculated with 10 mL starter culture and grown at 37°C, shaking at 170 rpm, until optical density (OD600) reached 0.4–0.6. Protein expression was then induced by addition of IPTG (Fluorochem) to a final concentration of 1 mM, as well as reduction of temperature to 18 °C and overnight incubation, shaken at 170 rpm. The following morning, the culture was harvested by centrifugation (4 000 rpm for 20 min), reconstituted in resuspension buffer (25 mM K·HEPES, pH 7.6, 500 mM KCl, 10% (v/v) glycerol) and stored at –80 °C prior to purification.

Thawed resuspensions were supplemented with 250 U BaseMuncher (Expedeon) and protease inhibitor cocktail (Roche) before cell lysis by sonication using a microprobe tip set to 35% amplitude, for 2 min total and with 10 s pauses between 5 s pulses. Lysate was clarified by centrifugation at 30 400 ×*g* for 40 min at 4 °C, and the supernatant immediately filtered through a 0.22 µm syringe filter. Filtered supernatant was flowed through a 5 mL HisTrap HP column (Cytiva) pre-equilibrated with resuspension buffer at 0.5 mL/min flow rate. After loading, the column with bound protein was washed with 5 column volumes (CV) of HisTrap buffer containing 20 mM imidazole (25 mM K·HEPES, pH 7.6, 500 mM KCl, 10% (v/v) glycerol, 1 mM TCEP, 20 mM imidazole), which was sufficient to restore a flat A280 baseline. This wash was then repeated using HisTrap buffer with 50 mM imidazole. A single-step elution with 3 CV of HisTrap buffer containing 250 mM imidazole was used after washing. Eluted proteins were concentrated in a 10 kDa molecular weight cutoff (MWCO) centrifugal filter (Amicon, Merck).

For SpyCatcher003, prior to its use for direct functionalisation of derivatised grids, concentrated protein was supplemented with 100 U TEV-His7 protease in 1X TEV protease reaction buffer (NEB) and incubated at 30 °C with agitation for 3 hr to cleave the His6-tag. The TEV-cleaved protein was then buffer exchanged using a 30 kDa MWCO centrifugal filter (Amicon, Merck) into buffer containing no imidazole and 50 mM KCl (25 mM K·HEPES, pH 7.6, 1 mM EDTA, 1 mM TCEP, 10% (v/v) glycerol, 50 mM KCl) and re-loaded onto the HisTrap column (Cytiva) pre-equilibrated with the same buffer. The column with bound sample was washed with 2 CV of the same buffer at a flow rate of 0.5 mL/min, and the flow-through, containing untagged SpyCatcher003 or mGreenLantern-SpyTag003, collected and concentrated as before.

Lastly, after concentration, mGreenLantern and SpyCatcher003 were injected onto a Superdex 200 size exclusion chromatography column (Cytiva) pre-equilibrated with size exclusion buffer (25 mM K·HEPES, pH 7.6, 200 mM KCl, 1 mM EDTA, 1 mM TCEP) at a flow rate of 0.25 mL/min. Peak fractions were visualised by SDS-PAGE and concentrated to 0.5 mg/mL before snap-freezing in liquid nitrogen and storage at –80 °C until use. The identity and purity of both SpyCatcher003 and mGreenLantern was verified by MS (BSRC Mass Spectrometry Facility, University of St Andrews).

### Preparation of functionalised GrOx-based affinity grids

Quantifoil R 2/1 300 mesh gold grids (Jena Bioscience) were washed sequentially with ddH2O and ethyl acetate, followed by plasma-cleaning (Harrick Plasma) at low RF (7.2 W) for 30 s. Graphene oxide dispersion (Sigma Aldrich) was diluted to 0.024 mg/mL in ddH2O and vortexed heavily for 1 min, before being applied to the plasma-cleaned grids as per established protocols (de Martín Garrido *et al*, 2021b). After GrOx application and air-drying, grids were incubated under vacuum for 30 min to maximise support film adherence.

The coupling of PEG compounds was carried out according to the method of Wang *et al* (2020) with minor modifications. DBCO-PEG4-NH2 (Jena Bioscience) stock was prepared at a final concentration of 100 mM in DMSO, and N3-PEG5k-NHS ester (Nanocs) stock was prepared at a final concentration of 40 mM in DMSO. To functionalise GrOx-coated grids, grids were immersed individually in 25 µL of 10 mM DBCO-PEG4-amine in 1.5 mL Eppendorf tubes and incubated on the bench overnight at room temperature. The following day, grids were washed 3× sequentially in 30 µL DMSO droplets, side-blotting the grid between each wash step. Grids were subsequently immersed in 25 µL of 4 mM N3-PEG5k-NHS ester in 1.5 mL Eppendorf tubes overnight at room temperature. Grids were washed the following day using the same procedure and, when necessary, stored in DMSO until use. When grids were to be used, they were washed a further 3× sequentially with ddH2O before air-drying.

To derivatise grids with SpyCatcher003, grids were washed as above and subsequently incubated on a 55 µL droplet of 1.5 µM SpyCatcher003 solution in one well of a 3D-printed grid floatation block (de Martín Garrido *et al*, 2021a), diluted from snap-frozen aliquots using size exclusion buffer, for 30 min. After this incubation, the grids were then transferred onto a 55 µL droplet in another well of a 3D-printed grid floatation block containing Tris buffered saline (TBS; 50 mM Tris·HCl, pH 8.0, 150 mM NaCl) for at least 30 min to quench any remaining NHS ester functional groups.

### Design and operation of a 3D-printable sample/grid flow-cell

Flow-cells (1-cell and 4-cell) were designed using Fusion 360 (Autodesk) and 3D-printed using Clear V4 resin on a Form 3 stereolithography 3D printer (Formlabs) housed at the Imperial College Advanced Hackspace (Imperial College London). The resolution of each print was set to 0.025 mm, which increased the printing time but ensured that the fine geometry of the flow-cells would be faithfully reconstructed. Due to the requirement of having threads be printed along the *z* (i.e. vertical) axis, the flow-cell device design had to incorporate threads at a 45° angle to the relevant internal geometry and flow-cells were printed using rafts with 0.5 mm touchpoints auto-generated in PreForm (Formlabs).

Inlet and outlet screw ports were designed at the back of the flow-cell, which interface seamlessly with ÄKTA Fingertight Connectors 1/16” (Cytiva) which can receive PEEK Tubing (internal diameter 0.75 mm, outer diameter 1/16”, Cytiva). The flow-cell was then connected via two circuits to a peristaltic pump (Gilson) using modified sections of PVC tubing (Gilson) connected at either end to a short length of 1/16” PEEK tubing. In the direction of peristalsis, the input circuit pumps sample or buffer from a reservoir into the inlet port of the device; the outlet circuit pumps from the device into a waste container. When operating the flow-cell, the tightness of the outlet circuit was manually adjusted each time to run slightly faster than the inlet circuit, to ensure that the grid wells did not over-fill. The 4-cell was designed by placing 4 completely independent flow circuits side-by-side.

### CNC milling

Toolpaths for CNC fabrication of 96-well plate designs were generated in Fusion 360 (Autodesk). Blocks were machined from 6 mm PTFE sheet using carbide tooling in a Haas CNC Mini Mill (HAAS Automation) housed at the Imperial College Advanced Hackspace (Imperial College London).

### On-grid affinity capture of MS2-SpyTag003

MS2-SpyTag003 and MS2-WT was produced according to established protocols (de Martín Garrido *et al*, 2020). Briefly, phage constructs were expressed overnight in Rosetta2 (DE3) pLysS cells under IPTG induction and the culture resuspended in buffer containing 50 mM Tris, pH 8.0, 5 mM MgCl2, 5 mM CaCl2, 100 mM NaCl, and further supplemented with 700 U RNAse A (Qiagen), 2500 U BaseMuncher (Expedeon) and 200 U Turbo DNAse (Thermo Fisher Scientific). Cells were lysed by sonication (50% amplitude, for 2 min total and with 30 s pauses between 30 s pulses) at 4 °C. Lysates were clarified by centrifugation at 16 602 ×*g* at 4 °C for 15 min and subsequently filtered through a 5 µm syringe filter (de Martín Garrido *et al*, 2020).

To perform the affinity capture, the 1-cell device was flushed with water followed by flow-cell washing buffer (25 mM K·HEPES, pH 7.6, 500 mM KCl, 1 mM EDTA, 0.096 mM DDM [0.8× CMC]) at approximately 1 mL/min using a peristaltic pump; the inlet and outlet pump circuits were manually adjusted until a steady, stable flow rate through the flow-cell was achieved and the out-flow operated at a slightly higher rate than the in-flow. Affinity grids were placed within the grid aperture during the buffer wash, before 10-fold diluted lysate (with an approximate concentration of 10 mg/mL) was flowed through the flow-cell at 0.5 mL/min for 10 mins. Immediately after the lysate, flow-cell washing buffer was flowed through the device at 1 mL/min for 25 mins. Grids were then lifted from the aperture using negative-action tweezers (Dumont) and negatively stained. For the comparison of specific MS2-SpyTag003 capture with the nonspecifically binding MS2-WT sample, grids were prepared in triplicate.

Affinity capture using the 4-cell for cryo-EM was performed similarly to the 1-cell, but with neat lysate, a lysate loading flow rate of 1 mL/min and up to 1 hr of both lysate loading and washing with flow-cell washing buffer.

### Fluorescence microscopy

Air-dried grids were placed GrOx-side-up on a glass slide pre-cleaned with 70% EtOH and mounted beneath a coverslip in ProLong™ Gold or ProLong™ Diamond Antifade Mountant gel (Thermo Fisher Scientific).

Images were acquired using an Axio Observer 7 widefield inverted microscope (Zeiss) fitted with a sCMOS Orca Flash 4 camera (Hamamatsu) and a Plan-Apochromat 100×/1.40 oil immersion lens (Zeiss). eGFP and mGreenLantern were detected using a 450/90 nm excitation filter and 500/50 nm emission filter. Full-frame images (1024 × 1024 pixels) were acquired in *z* steps of 2 µm to a total thickness of 16 µm, extending above and below the focal plane of each grid. Exposure times for each channel was kept consistent across comparative grids.

Images were processed with a combination of ZEN lite (Zeiss) and FIJI (Schindelin *et al*, 2012) software. For intensity analyses, a square region of interest (ROI) was drawn around each grid square. Fluorescence intensity maxima were plotted per slice and used to denote surface GrOx layers where derivatisation or functionalisation had occurred. Mean fluorescence intensity values were extracted for comparison from within the ROI for slices displaying maximal fluorescence intensity. For comparison where no appreciable fluorescence maxima existed (i.e. in any negative control grids), the brightfield images were used to visually assess stack depth and compare fluorescence accordingly.

### Electron microscopy

To examine GrOx coverage, GrOx-coated and -functionalised grids were imaged without further preparation on a Tecnai T12 microscope (Thermo Fisher Scientific) operated at 120 kV at room temperature (Imperial College, Electron Microscopy Facility). For negatively stained EM, prepared grids i.e. from affinity capture experiments using the flow-cell were held in a pair of negative-action tweezers and 5 µL 2% (w/v) UAc was added to the GrOx-coated side of the grid for 5 min. After this incubation, the UAc was blotted away using filter paper, and grids were supplemented with a further 1.5 µL of UAc, which was left to air-dry. These grids were then imaged on a Tecnai T12 microscope as above.

For the comparison of affinity capture with MS2-SpyTag003 against nonspecifically binding MS2-WT sample, we derived a quantification method from a previous study investigating affinity-purified viruses (Kiss *et al*, 2014). Across 3 independently prepared grids, 75 micrographs were collected for each of the MS2-WT and MS2-SpyTag003 samples on a Tecnai T12 electron microscope (FEI, Thermo Fisher Scientific) at a nominal magnification of 26 000× and an acceleration voltage of 120 kV. All particles in the field of view per micrograph were semi-automatically selected using BOXER (Tang *et al*, 2007); particles were characterised as viruses if they matched known morphological characteristics for MS2 as previously observed in-house (de Martín Garrido *et al*, 2020). Statistical comparison of mean particle counts across triplicate grids was performed by unpaired *t*-test (GraphPad Prism 9).

To prepare affinity grids for cryo-EM data collection, grids prepared using the flow-cell were stored on TBS buffer after affinity capture until vitrification. Affinity grids were transferred to the chamber of a Vitrobot Mark IV (FEI, Thermo Fisher Scientific) at 20 °C and 100% humidity, where the sample side of the grid was supplemented by the addition of 4 µL TBS buffer before being blotted for between 4–6 s with a blot force of 0 and subsequent plunge freezing into liquid ethane. We collected micrographs of MS2-SpyTag003 sample on a Glacios electron microscope (FEI, Thermo Fisher Scientific) at a nominal magnification of 63 000× and an acceleration voltage of 200 kV.

#### Abbreviations

2D/3D: 2/3-Dimensional
(Cryo-)EM: (Cryogenic) Electron Microscopy
DBCO: Dibenzocyclooctyne
DMSO: Dimethyl Sulfoxide
FITC: Fluorescein Isothiocyanate
GrOx: Graphene Oxide
NHS: N-hydroxy Succinimide
NS: Negative Staining
PEG: Polyethyleneglycol

## Acknowledgements

The authors would also like to thank all the members of the Section for Structural and Synthetic Biology at Imperial College London who have helped test these techniques, and Paul Simpson at the Centre for Structural Biology. This work utilised expertise and prototyping equipment at the Imperial College Advanced Hackspace, microscopy equipment at the Facility for Imaging by Light Microscopy (FILM) at Imperial College London and electron microscopy expertise and equipment within the Imperial College Centre for Structural Biology.

## Funding

This work was funded by the Wellcome Trust and the Royal Society through Sir Henry Dale Fellowships (206212/Z/17/Z and 206212/Z/17/A) to CHSA and (211230/Z/18/Z) to DTR, and by the UK Engineering and Physical Sciences Research Council (EPSRC) through award EP/R014000/1 to PSF. The Imperial College Advanced Hackspace is supported by the Higher Education Innovation Fund.

## Conflict of interest statement

The authors declare that they know of no conflicts of interest with respect to this work.

## Notes

### Competing Interest Statement

The authors have declared no competing interest.

### Summary of Updates

The manuscript has been revised to include improvements and additions to the described equipment as well as data on it's use in the generation of cryo-EM samples and images.

